# A retinotopic reference frame for space throughout human visual cortex

**DOI:** 10.1101/2024.02.05.578862

**Authors:** Martin Szinte, Gilles de Hollander, Marco Aqil, Inês Veríssimo, Serge Dumoulin, Tomas Knapen

## Abstract

We perceive the world as stable despite our rapid eye movements. To explain our sense of visual stability, it has been suggested that the brain encodes the location of attended visual stimuli in an external, or spatiotopic, reference frame. However, such spatiotopy is seemingly at odds with the fundamental retinotopic organization of visual inputs. Here, we probe the spatial reference frame of vision using ultra-high-field (7T) fMRI and voxel-level receptive field modeling, while manipulating both gaze direction and spatial attention. To manipulate spatial attention, participants performed an equally demanding visual task on either a bar stimulus that traversed the visual field, or a small stimulus at fixation. To dissociate retinal stimulus position from its real-world position the entire stimulus array was placed at one of three distinct horizontal screen positions in each run. We found that population receptive fields in all cortical visual field maps are pinioned to the retina, irrespective of how spatial attention is deployed. This pattern of results is strong evidence for a fully retinotopic reference frame for visual-spatial processing. Reasoning that a spatiotopic reference frame could independently be computed at the population level of entire visual areas rather than in individual voxels, we additionally used Bayesian decoding of stimulus location from the BOLD response patterns in visual areas. We found that decoded stimulus locations also adhere to the retinotopic frame of reference. Again, this result holds for all visual areas and irrespective of the deployment of spatial attention. Our findings reorient the search for visual stability mechanisms toward transient sensorimotor interactions rather than static spatiotopic maps.

## Introduction

Eye movements rapidly alter the projection of visual objects on the retina. Nevertheless, our subjective visual perception is stable and invariant to these eye movements. This “*space constancy*” phenomenon presents researchers with a conundrum: how do our brains build a stable impression of the world based on such fleeting, jumbled sensory inputs? One major step to understanding space constancy would be to answer the question of how the information of visual objects’ location is represented by the visual system.

Visual location is first encoded by the retina^*1*^, whose topography is maintained in subsequent processing stages^*2,3*^. This fundamental “retinotopic” reference frame^*4,5*^ maintains the retinal photoreceptor organization, and neural responses in this frame are modulated by gaze direction^*6,7*^. This combination of action-related and visual sources of information is thought to allow the brain to localize objects in the outside world^*8–10*^. But the precise mechanisms underpinning world-centered localization, including the role of spatial attention, remain a matter of debate^*11–14*^. Specifically, an open fundamental question is whether gaze modulation of visual information leads to the encoding of locations of visual stimuli in a spatiotopic (i.e. head- or world-centered) frame of reference.

We reasoned that questions of spatial encoding are best answered using an experimentation and analysis paradigm that provides detailed, robust estimates of spatial tuning at the single-voxel level. Thus, we performed population receptive field (pRF) mapping^*15*^ at ultra-high-field (7T) whole brain fMRI to probe the spatial reference frame of vision. This approach allows us to focus at the level of local neural populations sampled by individual voxels without having to average across regions. We determined voxels’ pRF position while participants directed their gaze to three distinct locations, allowing us to test whether pRFs are pinioned to the outside world (the *spatiotopic* hypothesis), or fixed to the retina (the *retinotopic* hypothesis). To factorially study the role of spatial attention, participants performed an equally demanding orientation-discrimination task either at fixation or on a moving visual-mapping stimulus.

We found that throughout the cortical visual hierarchy, pRFs systematically shift with gaze, being fixed to the retina irrespective of where spatial attention is deployed. More informative voxels regarding our hypotheses showed a tendency to be more retinotopic. Next, to probe the visual reference frame at the level of entire visual field maps, we adapted a Bayesian decoding method^*16–18*^ to infer the spatial location of our visual stimuli from the pattern of voxel responses in entire visual areas. In line with our single-voxel results, we found that decoded stimulus locations are fixed to the retina, again favoring the retinotopic hypothesis. Altogether our results suggest that location is represented throughout the visual hierarchy by populations of receptive fields operating in a retinotopic reference frame.

## Results

### PRF modeling and attention effect

Participants were instructed to continuously fixate a bull’s-eye on a black background while horizontal and vertical bars swiped across the full extent of the screen (Fig. 1a, *full screen*). They reported the orientation of filtered noise patterns presented either within the moving bar (attend-bar condition) or within the fixation bull’s-eye (attend-fix condition). We titrated the difficulty of the task with a staircase procedure which adjusted the orientation dispersion coefficient of pink noise patterns inside the presented stimuli (Fig. 1b). Participants maintained the expected staircase performance level (Fig. 1c, *full screen*, attend-bar: 74.06% ± 2.50; *full screen*, attend-fix: 80.26% ± 0.40), confirming their ability to attend to the noise patterns presented in either the periphery or at fixation with equal performance. To ensure correct fixation, participants were trained outside the scanner and fixation performance was checked in the scanner using DeepMREye^19^ (see Methods). Apart from the attention task, the stimulus presentation sequence of *full screen* runs was identical (Fig. 1d, top), allowing us to average signal timeseries across the attention tasks (Fig. 1e) into a single, high-SNR fiducial timeseries. From each voxel’s fiducial timeseries, we quantified the screen locations at which a stimulus evokes a visual response by estimating its population receptive field (pRF) properties. Specifically, we fit a linear, isotropic Gaussian pRF model to find optimal location (pRFx and pRFy) and size (pRFsize) parameters that quantify the spatial tuning of the neural population within each voxel. Fig. 1f shows an example of a V1 voxel fiducial timeseries and its best-fitting pRF model. From the position parameters, we derived pRF eccentricity and pRF angle maps that can be projected onto individual participants’ cortical surfaces (Fig. 1g) to delineate visual field maps as regions of interests (see Methods). We next repeated this analysis using timeseries of attend-bar and attend-fix runs averaged separately (Fig. 1h, top), using the fiducial timeseries outcomes as starting parameters. Fig. 1h show an inflated cortex of a participant displaying the change in explained variance (R^2^) resulting from manipulating spatial attention. Using the 250 best-fitting voxels per ROI (see Methods), we found a significant improvement of the explained variance when comparing the attend-bar to the attend-fix conditions for all ROIs (attend-bar: 0.51 > R^2^ > 0.18 vs. attend-fix: 0.43 > R^2^ > 0.15, 0.0117 > ps > 0.0001, two-sided p values) except area LO (attend-fix: R^2^: 0.38 ± 0.03 vs. attend-bar R^2^: 0.36 ± 0.04, two-sided p = 0.24). This pattern of results was similar when including all voxels contained within the ROIs. Taken together, behavioral and pRF mapping results illustrate classical effects of visual spatial attention: an improvement of orientation discrimination abilities^20,21^ correlated with an improvement of the corresponding fMRI BOLD signal quality^22,23^. Our design yielded reliable voxel-wise pRF estimates under balanced attentional demands, ensuring that any reference-frame effects could not be attributed to task differences.

**Fig. 1.**
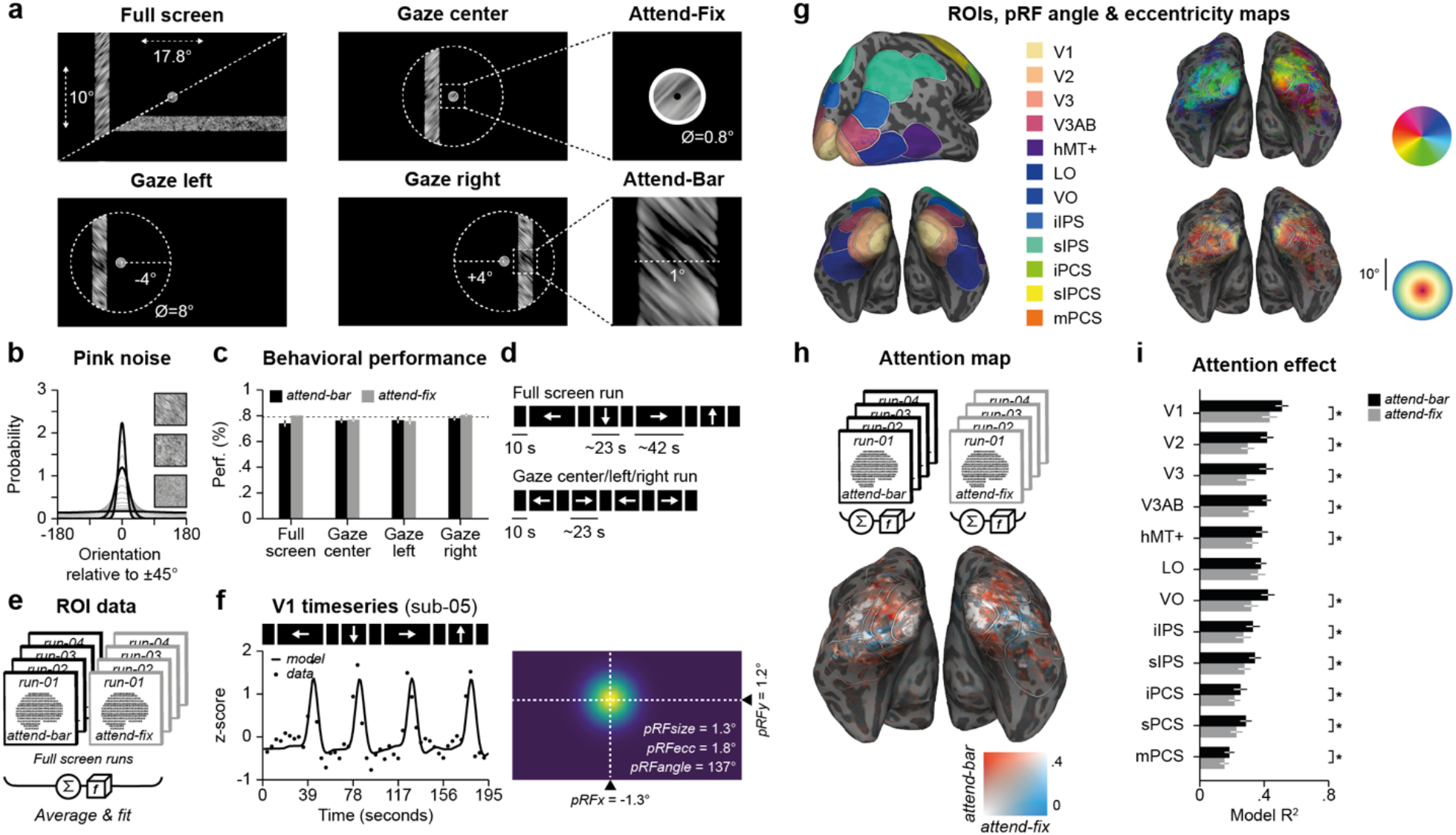
Methods, pRF modeling and attention effect. **a**. Participants fixated a bull’s-eye in either *full screen* (top left panel), *gaze center* (top center), *gaze left* (bottom left) and *gaze right* runs (bottom center). They reported the orientation of pink noise patterns (+45° or –45°) presented either within the fixation bull’s-eye (*attend-fix*) or within the bar (*attend-bar*). **b**. We titrated the difficulty of the orientation discrimination task by varying the dispersion coefficient of the orientation filter applied to the pink noise pattern. The figure presents probability distributions of orientations contained within the pink noise patterns as a function of the dispersion coefficient. Right insets showcase three levels of difficulty from non-oriented random noise (bottom) to the most oriented pattern used (top). **c**. Group performance (n = 8) obtained in the *attend-bar* (black) and *attend-fix* (gray) conditions. Error bars show ±SEM and the dashed line shows the staircase convergence level (∼79% correct). **d**. In the *full screen* and *gaze center/left/right* conditions, a bar moved in different directions (black rectangles with arrows) interleaved with periods in which only the fixation bull’s-eye was shown (empty black rectangles). **e**. To determine regions of interest (ROIs), we averaged *full screen* runs and fit a linear pRF model. **f**. Example V1 timeseries and its best explained pRF model and parameters. **g**. Single participant (*sub-03*) ROIs and pRF angle and eccentricity maps projected on his inflated brain. Participants brains visualizations are available online (invibe.nohost.me/gazeprf). **h**. Map of the effect of spatial attention obtained by comparing the explained variance of *attend-bar* and *attend-fix* conditions separately. Reddish colors of the inflated cortex illustrate the effect of spatial attention. **I**. Model explained variance (R^2^) observed for the best-fitting voxels of each ROI in the *attend-bar* (black) and *attend-fix* (gray) conditions. Error bars show ±SEM, asterisks show significant difference between conditions (two-sided *p* < 0.05).

### Out-of-sample predictions show retinotopic encoding of visual space

To test the spatial reference frame, we used pRFs from the full-screen condition to predict responses when gaze shifted left, right, or center (Fig. 1a). The entire mapping stimulus sequence was centered on and identical relative to these fixation locations. Bar travel was restricted to vertical bars moving rightwards or leftwards, vignetted by an invisible circular aperture (Fig. 1a and Fig. 1d) in order to efficiently sample the dimension in which changes in spatial sampling are expected. As in the *full screen* runs, participants reported the orientation of noise patterns presented within the bar (attend-bar) or within the fixation bull’s-eye (attend-fix). They reached the staircase expected performance level in the *gaze center* (Fig. 1c, attend-bar: 76.27% ± 1.67; attend-fix: 76.95% ± 1.48), the *gaze left* (Fig. 1c, attend-bar: 76.59% ± 2.30; attend-fix: 75.88% ± 2.49) and *gaze right* runs (Fig. 1c, attend-bar: 78.09% ± 1.66; attend-fix: 80.61% ± 1.25). Using the *full screen* runs’ pRF parameters, we calculated predicted timeseries in the retinotopic and spatiotopic reference frame (Fig. 2a). Specifically, to obtain retinotopic predictions we added the gaze direction change to the pRFx parameter (x-coordinate of the pRF) obtained in the corresponding *full screen* runs (Fig. 2b, retinotopic prediction). The spatiotopic hypothesis supposed that pRFx does not vary as a function of the gaze directions (Fig. 2c, spatiotopic prediction). No estimation of spatial pRF parameters was performed at this stage. Fig. 2b-c illustrate the predicted and measured timeseries of V1 and hMT+ voxels. We highlighted these regions to relate them to former studies reporting spatiotopic effects^11,12^, and to highlight the effects of different pRF sizes on the BOLD signal timeseries. Importantly, the *gaze center* condition here can serve as a baseline: the spatiotopic and retinotopic predictions necessarily produce identical pRFx parameters. We computed the change in the pRF model explained variance (R^2^) between the *gaze left* and *gaze right* runs as compared to the *gaze center* runs. We found, again using the best 250 voxels across ROIs, significantly lower explained variance for the spatiotopic relative to the retinotopic prediction in both the attend-bar (Fig. 2d, spatiotopic prediction: +2.36% > R^2^ change > -28.91% vs. retinotopic prediction: +3.96 > R^2^ change > +1.09%, 0.0195 > ps > 0.0001, two-sided p values) and attend-fix conditions (Fig. 2e, spatiotopic prediction: +4.61% > R^2^ change > -23.59% vs. retinotopic prediction: +6.14 > R^2^ change > -1.12%, 0.0352 > ps > 0.0001, two-sided p values). Similar effects were found when including all voxels from the ROIs. Across visual areas, retinotopic predictions consistently explained more variance than spatiotopic predictions, in both attention conditions. This indicates that voxel responses tracked retinal rather than world-centered spatiotopic coordinates.

**Fig. 2.**
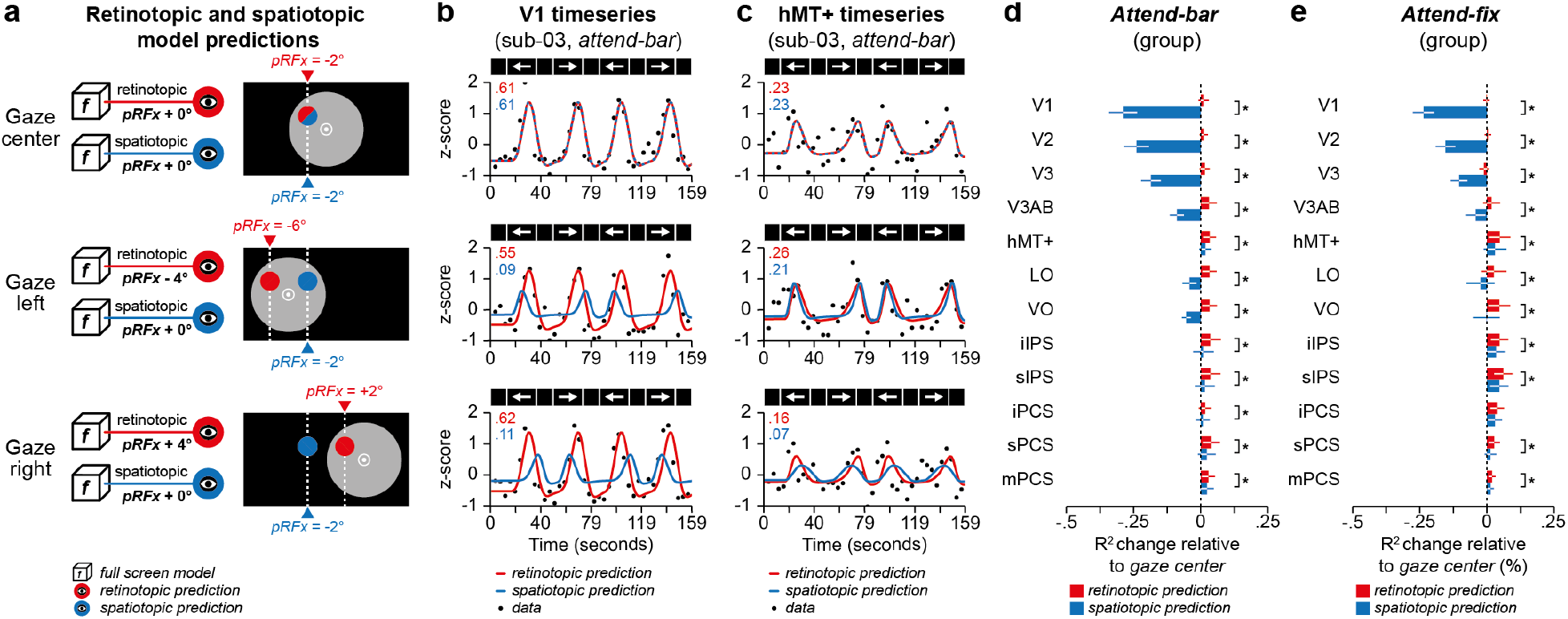
Out-of-sample retinotopic and spatiotopic predictions. **a**. Retinotopic predictions are obtained by adding the change in gaze direction to the *full screen pRFx* parameters (e.g., *pRFx*_*gaze left*_ = *pRFx*_*full screen*_ *– 4°*). The spatiotopic hypothesis dictates that the *pRFx* should stay identical irrespective of the gaze direction (i.e., *pRFx*_*gaze left*_ = *pRFx*_*gaze right*_ = *pRFx*_*gaze center*_ = *pRFx*_*full screen*_). **b-c**. Example retinotopic (red) and spatiotopic (blue) timeseries predictions for V1 (b) and hMT+ voxels (c) in the *gaze center* (top row), *gaze left* (middle row) and *gaze right (bottom row) attend-bar* runs of one participant (sub-03). Leftmost inset values show corresponding model R^2^. **d-e**. Retinotopic (red) and spatiotopic predictions (blue) change in explained variance between *gaze left/right* and *gaze center* conditions across ROIs. Note that only spatiotopic prediction of V1, V2 and V3 voxels display a significant change as compared to zero in both attention conditions (V1/V2/V3: 0.0156 > *ps* > 0.0001; two-sided *p* values), as well as V3AB (two-sided *p* = 0.0156) and VO in the attend-bar condition (two-sided *p* = 0.0078). Error bars show ±SEM, asterisks show significant difference between retinotopic and spatiotopic predictions (two-sided *p* < 0.05).

### Fitting pRFs under gaze shifts

We next asked whether receptive fields shifted with gaze. Fitting pRFs separately for each gaze as a function of attention condition revealed that receptive field centers consistently translated with eye position, as predicted by retinotopy, rather than remaining fixed on the screen. We now fit the horizontal pRF position parameter in the gaze conditions while keeping the other pRF parameters constant (pRFy and pRFsize). To avoid any bias due to the starting points of the fitting procedure, we initiated parameters from distinct horizontal pRFx position values referenced against either the gaze position (Fig. 3a, retinotopic starting points) or the screen center (Fig 3a, spatiotopic starting points). In a subsequent stage, we optimized the horizontal pRF position parameter starting from the highest explained variance model (R^2^) of the first stage. We formulated two distinct predictions based on our two competing hypotheses. As before, the *gaze center* condition is a reference for both hypotheses, since they produce identical predictions. In the retinotopic reference frame hypothesis, horizontal pRF position in the *gaze left* and *gaze right* conditions shifts based on gaze direction (Fig. 3b, retinotopic prediction). Conversely, in the spatiotopic reference frame hypothesis, horizontal pRF position shouldn’t differ from that observed when participants look straight ahead (Fig. 3b, spatiotopic prediction). Figs. 3c & 3d show these comparisons across participants for best-fitting voxels of V1 and hMT+ using data modeled from the attend-bar and attend-fix runs respectively. In accordance with previous studies^11–13^, pRF x positions follow the retinotopic prediction also when this parameter is directly estimated from the data, irrespective of where participant deploy their spatial attention (Fig. 3c-d, top). These results suggest that both V1 and hMT+ use a retinotopic framework, regardless of spatial attention’s focus (Fig. 3c-d, bottom).

**Fig. 3.**
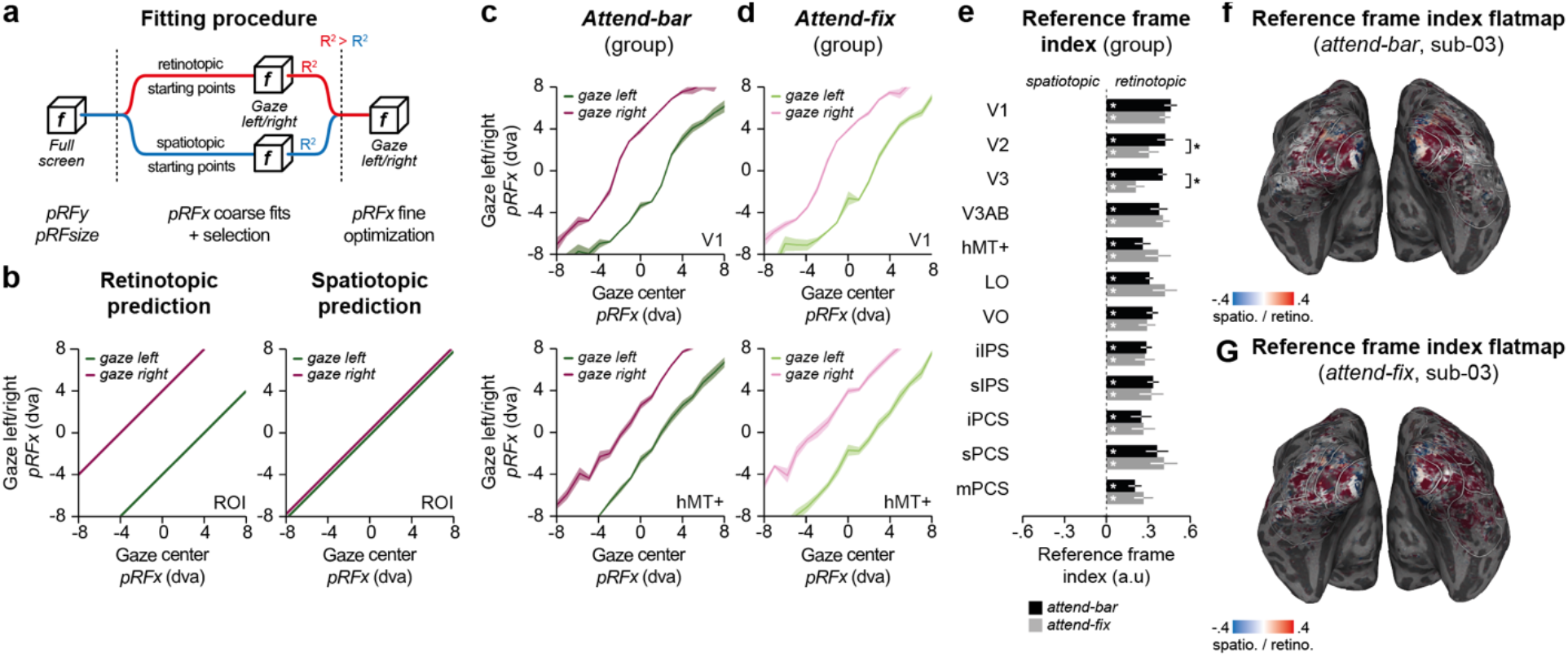
Fitting retinotopic and spatiotopic pRF models. **a**. Fitting procedure. We used a coarse-to-fine optimization fitting procedure in which we kept the *pRFy* and *pRFsize* parameters from the corresponding *full screen* runs. In order to avoid bias between the spatiotopic and retinotopic model, we first used two different sets of parameter starting points. These starting points were based on either the retinotopic (red) or the spatiotopic (blue) hypotheses. For a next optimization stage, we selected the parameters producing the highest fit quality. **b**. Retinotopic and spatiotopic predictions. The *gaze center pRFx* as a function of the *gaze left/right pRFx* will either shift (left panel, retinotopic prediction) or remain at the same position (right panel, spatiotopic prediction) when comparing the *gaze center* with the *gaze left* (green) or *gaze right* conditions (purple). **c-d**. Each panel shows the group average of 16 equal bins of the *pRFx* obtained in the *gaze center* runs as a function of the corresponding group averaged bins of the *pRFx* obtained in the *gaze left* (purple) and *gaze right* runs (green) for the *attend-bar* (c, dark colors) and *attend-fix* conditions (d, light colors). Error areas show ±SEM. **e**. Group average reference frame index (RFI) observed in the *attend-bar* (black) and *attend-fix* conditions (gray) for the best-fitting voxels of each ROI. Error bars show ±SEM, white asterisks show significant RFI as compared to a null effect (two-sided *p* < 0.05). Black asterisks indicate a significant difference between the *attend-bar* and *attend-fix* conditions (two-sided *p* < 0.05). **f-g**. Reference frame index maps obtained by projecting the RFI obtained in the *attend-bar* (f) and *attend-fix* conditions (g) on inflated cortex using a unimodal color scale (blue: spatiotopic vs. red: retinotopic) for a participant (sub-05, same as in Fig. 1).

To evaluate quantitatively which reference frame best explains our results at the level of individual voxels across ROIs, we computed a reference frame index (RFI, see Methods), as in previous studies^11,13^. RFI quantifies the difference in explained variance between the two reference frame models on a scale between -1 and 1, where pure spatiotopy corresponds to -1, pure retinotopy to +1, and noise results in an RFI value of 0. Fig. 3e shows RFI obtained with the best-fitting voxels of each ROI for the attend-bar and attend-fix conditions. Across participants, RFI were significantly positive for all ROIs in the attend-bar (0.46 > RFI > 0.20, 0.0312 > ps > 0.0001, two-sided p values) and attend-fix conditions (0.42 > RFI > 0.21, 0.0312 > ps > 0.0001, two-sided p values). Moreover, RFI values did not differ between conditions in any ROIs (attend-bar: 0.93 > ps > 0.38, two-sided p values) other than V2 (attend-bar RFI: 0.42 ±0.06 vs. attend-fix RFI: 0.30 ± 0.08, two-sided p < 0.01) and V3 (attend-bar RFI: 0.40 ±0.03 vs. attend-fix RFI: 0.21 ± 0.07, two-sided p < 0.01). Similar statistics were observed when including all voxels. This analysis again suggests that at the level of individual voxels, visual location is encoded in a retinotopic reference frame irrespective of the deployment of attention. Fig. 3f and 3g illustrate these effects by projecting the RFI onto the cortical surface of a single participant when they attended stimuli presented within the bar (Fig. 3f) or the fixation bull’s-eye (Fig. 3g).

To ensure the robustness and validity of these results, we identified two dimensions that should make a voxel a reliable and unbiased indicator of spatial reference frame: their spatial location, and their signal fidelity. Regarding the first, spatial dimension, a natural and neutral quantification is to what extent voxels’ pRFs overlap with the stimulus aperture in the center condition, when spatiotopic and retinotopic predictions are identical. The second dimension of signal fidelity reflects our ability to use a voxel’s responses for our inference at all. We expect voxels with low explained variance in the full screen runs to be unreliable indicators of spatial reference frame. Their fiducial spatial parameters are more uncertain, and in addition to having an unstable baseline for quantification, their estimated pRFx parameters in the gaze conditions are likely to suffer from the same lack of signal fidelity. We binned voxels’ RFI values, both according to their fiducial pRF’s model overlap with the *gaze center* stimulus aperture, as well as their within-set explained variance ratio in the fiducial pRF mapping timeseries. In this visualization (Fig. 4), we expect the reliable and spatially informative indicators to live in the upper-right quadrant. Interestingly, for all ROIs this upper-right quadrant has RFIs most strongly indicating the retinotopic frame of reference, dropping off gradually when moving away from the best-fitting, most spatially informative voxels. Taken together, these results indicate that voxels with stronger inferential contributions show stronger evidence of a retinotopic reference frames.

**Fig. 4.**
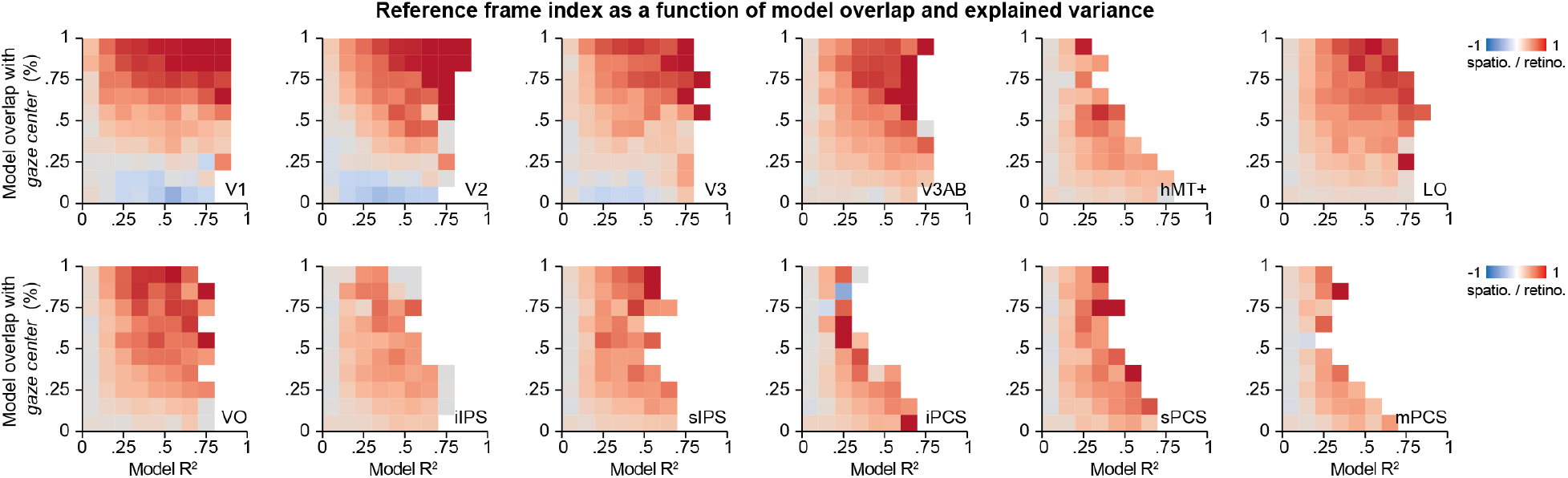
Reference frame index as a function of model overlap and explained variance. We ordered and binned voxels in two dimensions according to spatial relevance (vertical dimension, quantified as pRF overlap with the *gaze center* aperture) and explained variance (horizontal dimension). A strong gradient with higher RFI values emerges in the more informative voxels (upper right quadrant) across all visual ROIs, indicating that voxels with stronger inferential contributions show increasingly retinotopic reference frames.

### Bayesian decoding of stimulus positions under gaze shifts

Finally, we used a Bayesian decoder to reconstruct stimulus position from distributed responses, providing a population-level test of reference frame. We adapted a Bayesian decoding method developed for estimating, from a pattern of BOLD responses, a full posterior distribution along a single stimulus dimension, namely visual orientation^16^, to the multiple stimulus dimensions of visual space. Our novel method decodes spatial locations of the stimulus bar on a TR-by-TR basis (see Methods), based on pRF estimates and residual covariance structure from the *full screen* runs (Fig. 5a). Contrary to techniques which decode a single stimulus value most consistent with the observed brain data, Bayesian decoding also produces uncertainty estimates for the decoded features. This decoded uncertainty has been shown to relate to sensory uncertainty and decision confidence^16,24^. Our method determines, for every TR, the full posterior probability along spatial bar location parameters based on the per-TR BOLD pattern in the *gaze center, gaze left* or *gaze right* conditions (Fig. 5b). Because this posterior distribution lives in visual space it can be compared to the ground truth of what was actually shown to participants on the screen.

**Fig. 5.**
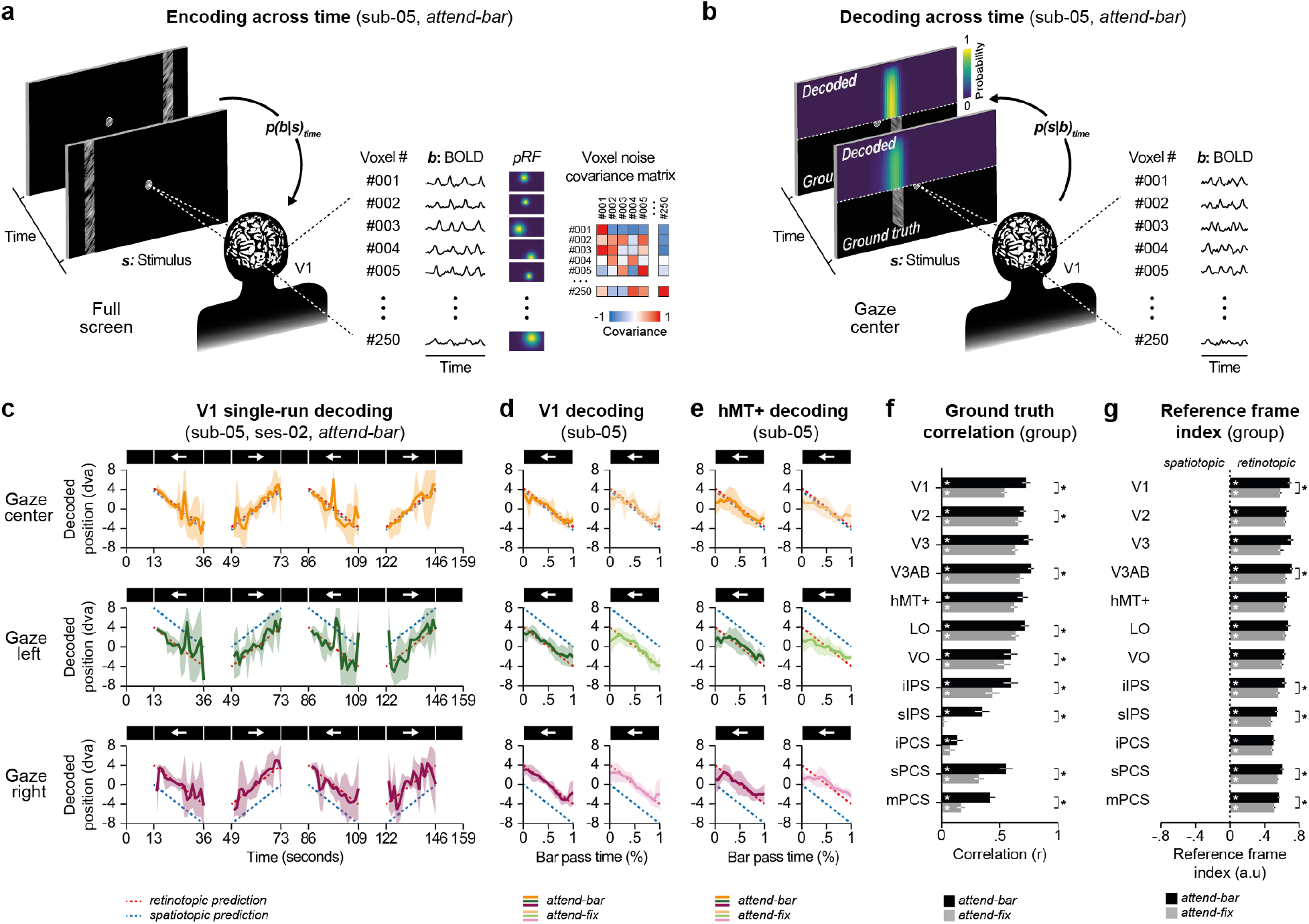
Bayesian decoding. **a**. Our encoding procedure consists of determining a likelihood *p(b*|*s)*_*time*_ defined as the probability of observing fMRI BOLD signal (*b*) as a function of the stimulus (*s*) presented in the *full screen* condition. For this analysis we use the spatial tuning and noise correlations of the 250 best-fitting voxels per ROI. **b**. The decoding procedure illustrated here computes the posterior probability over time *p(s*|*b)*_*time*_ using independent recorded BOLD signals for example from the *gaze center* condition. This procedure results in a direct comparison of decoded and actual bar position over time (see comparison of posterior distribution sample and ground truth). **c**. Decoded bar position as function of the time in the gaze center (top, orange), the gaze left (middle, green) and the gaze right (bottom, purple) conditions. Graphs present decoding from individual runs (from the second session) using V1 voxels activity (*attend-bar* condition) of a participant (sub-05). Predictions are illustrated as red and blue dashed lines for the retinotopic and spatiotopic predictions, respectively. Error areas show the posterior distribution STD. **d-e**. Average decoded bar position across bar passes and sessions for best-fitting voxels of V1 (d) and hMT+ (e) respectively, for *gaze center* (top), *gaze left* (middle) and *gaze right* conditions(bottom), in the *attend-bar* (left panels, dark colors) and *attend-fix* conditions (right panels, light colors). Error areas show ±SEM. **f**. Group averaged correlation between the decoded bar position and the ground truth. Conventions are as in Fig. 3e. **g**. Group average reference frame index observed in the *attend-bar* (black) and *attend-fix* conditions (gray) for the best-fitting voxels of each ROI. Conventions are as in Fig. 3e.

Crucially, we can define distinct spatiotopic and retinotopic predictions for the reference frame in which the decoded bar position is hypothesized to reside. As our decoder was trained on the *full screen* runs in which participants fixated straight ahead, the spatiotopic hypothesis predicts that our decoder will sense the change of gaze as a shift of the bar position in the opposite direction, as its position on the screen has changed. Conversely, the retinotopic hypothesis predicts that the decoded position will be centered on the coordinates of the *full screen* stimulus, irrespective of gaze direction. Fig. 5c shows the decoded position of the bar (i.e., the peak of the bar position distribution) as observed when considering V1 voxels from single runs of the *gaze center*, the *gaze left* and the *gaze right* conditions. Fig. 5d-e shows, for a single participant, the same result averaged across runs and bar passes, for V1 (Fig. 5d) and hMT+ voxels (Fig. 5e). To quantify decoding performance, we computed the correlation between the decoded position and the retinotopic ground truth of the moving bar (Fig. 5f). Correlations in the *attend-bar* condition were significantly positive for all ROIs (0.77 > r > 0.13, 0.0391 > *ps* > 0.0001, two-sided *p* values) with the strongest correlations for low-level visual areas (e.g., V1, r = 0.73 ± 0.04, *p* < 0.01; hMT+, r = 0.70 ± 0.05, *p* < 0.0001, two-sided *p* values). Correlations were significantly positive in the *attend-fix* condition across all ROIs (0.67 > r > 0.17, 0.0078 > *ps* > 0.0001, two-sided *p* values), except for sIPS (r = 0.02 ± 0.03, two-sided *p* = 0.49) and iPCS (r = 0.07 ± 0.05, two-sided *p* = 0.21). Furthermore, when participants directed spatial attention to the moving bar, correlations were significantly increased when compared to the *attend-fix* condition in all ROIs (*attend-bar* vs. *attend-fix*, 0.0391 > *ps* > 0.0001, two-sided *p* values), apart from V3 (two-sided *p =* 0.09), hMT+ (two-sided *p* = 0.06) and iPCS (two-sided *p* = 0.21).

To answer our central question concerning visual reference frames, we computed a decoding-based RFI analogous to the earlier RFIs (see Methods) for each participant and ROI (Fig. 5g, see *Methods*). Across participants, RFIs were significantly positive across all ROIs, indicating a predominantly retinotopic reference frame in both the *attend-bar* (0.71 > RFI > 0.51, 0.0078 > *ps* > 0.0001, two-sided *p* values) and *attend-fix* conditions (0.64 > RFI > 0.47, 0.0078 > *ps* > 0.0001, two-sided *p* values). Deployment of attention to the bar stimulus increased the RFI for V1, V3AB, iIPS, sPCS and mPCS (0.0391 > *ps* > 0.0078, two-sided *p* values), indicating that decoding fidelity increases the strength of retinotopic representations in both low and high levels of the visual hierarchy. Lastly, we inspected whether the decoder’s uncertainty perhaps provides residual indications for any role of spatiotopic spatial coding of visual locations. Specifically, the mismatch between train and test set due to differences in gaze location should increase uncertainty for *gaze left* and *gaze right* conditions relative to the *gaze center* condition. We quantified the uncertainty of the decoder by calculating the dispersion of the posterior along the horizontal position dimension. In the intermediate ROIs of the visual system, specifically in areas like V3AB, hMT+, LO, VO, and iIPS, we observed distinct outcomes of spatial attention. Specifically, when attention was directed towards the bar stimulus, there was a notable reduction in decoder uncertainty, aligning with the previously explained impacts on decoding fidelity (Supp. Fig. 1a). The impact of decoder uncertainty on attention was more pronounced than any discernable influence of gaze direction, which was negligable (Supp Fig. 1b).

Altogether decoded stimulus positions consistently aligned with retinal coordinates. Attention sharpened decoding fidelity but never altered the reference frame. Thus, at the population level as well, spatial coding remained strictly retinotopic.

## Discussion

Our goal was to determine whether human visual cortex encodes space in retinal or world-centered coordinates. Using ultra-high-field fMRI and quantitative pRF modeling, we factorially manipulated gaze and attention to directly pit retinotopic and spatiotopic hypotheses against each other. Across all visual field maps, receptive fields shifted with gaze, demonstrating strictly retinotopic encoding. Attention enhanced representational fidelity but did not alter the reference frame. Thus, visual cortex fundamentally encodes space in retinal coordinates, setting clear boundary conditions on proposed solutions to the space constancy problem. These results highlight dynamic motor–sensory interactions, such as remapping based on efference copy signals, as the more plausible route to stable perception. Several design choices ensured the robustness of this conclusion. Ultra-high-field fMRI provided the SNR necessary for voxel-level modeling, and our pipeline controlled for differences in signal quality that might otherwise bias results toward one reference frame^13^. Because peripheral visual transients can contaminate central responses^7,25^, we used static gaze positions within runs and limited peripheral stimulation, preventing artifactual spatiotopic patterns. Most importantly, we directly compared quantitative predictions of retinotopic versus spatiotopic models, both at the level of single voxels and of distributed responses. In each case, data favored retinotopy.

Across subsequent analyses these direct comparisons between data and predictions favored the retinotopic reference frame hypothesis as the fundamental spatial encoding scheme of the visual cortex. As for the role of attention, our results align with previous studies indicating that spatial attention enhances the fidelity of spatial representations^*22,23*^. Spatial attention is known to modulate pRF position in accordance with the predictions of gain-field models^*26,27*^. Our results indicate that enhanced signal fidelity afforded by spatial attention renders visual cortex more, not less (Figs 3e, 4f-g), retinotopically organized^*11*^. We conclude that the differential deployment of spatial attention does not result in changes in the retinotopic spatial reference frame of visual cortex. Also, voxels that are more informative due to their signal fidelity and/or ground-truth spatial tuning more strongly point to a retinotopic reference frame. Spatiotopic RFI values were consistently associated with worse data quality and more uninformative spatial preferences.

Every day, we experience a stable visual world across our eye movements: an experience far removed from the flutter of images projected onto our retinae. The present results reframe and highlight the fundamental question of space constancy across gaze changes, shifting the focus to mechanisms that achieve it with a robustly retinotopic visual cortex. Different solutions to the space constancy problem have been proposed^28,29^, we here briefly review these solutions through the lens of our results. A first category of solutions to the space constancy problem relies on the use of proprioceptive information of the eye position in the skull. An influential computational model^30^ supported by numerous findings in both low-level^31,32^ and high-level visual areas^33,34^ proposed that the proprioceptive signal of gaze direction could be used to recover stimulus position across eye movements. These findings suggest that the gain of retinotopically organized neurons is modulated by a field which encodes the position of the animal’s eye. It is known from electrophysiological studies that although neighboring neurons encode similar locations in retinotopic visual space, they experience wildly varying gain-field effects^32^, reflecting a salt-and-pepper organization. We did not find an influence of gaze position at the level of individual voxels nor at the level of visual areas, which is possibly in line with this organization at the single neuron level. Other than the potential absence of structured gain field maps, other reasons, however, might have prevented us from finding them. First, visual display size used in fMRI studies only allows gaze direction changes smaller than those used in non-human primates’ electrophysiology, although the magnitude of our gaze shifts is in the range of most saccades made during naturalistic viewing. Second, gain fields from animal studies involved modulation of neuronal activations while our results rely on fMRI BOLD signals, which only is an indirect proxy of neuronal firing rates^35,36^. Third, the timescales at which these eye position gain fields impact spatial processing may be faster than those of our experimental runs. Thus, our results are consistent with gain-field models in that they assume a fundamentally retinotopic substrate. What our findings rule out is an early transition to spatiotopic encoding in visual cortex: regardless of attention or gaze, responses aligned with retinal coordinates^37^.

A second category of solution involves the use of an efference copy^38^ which predict the sensory consequences of upcoming eye movements^39,40^. These corrections have been proposed to be performed on retinotopically organized visual maps with the process being orchestrated by visual attention^41,42^. Remapping has been observed at the single neuron level in distinct visual maps^43^, fMRI studies^44–46^ and human behavior^47,48^. Our results are compatible with efference-copy mechanisms: they indicate that spatial coding remains retinotopic, creating the substrate upon which predictive updating must act. It is important to note that in order to model voxel activity we asked participants to maintain steady gaze throughout each run. They change gaze directions only between runs, and not across trials, an explicit design choice to avoid any peripheral visual stimulation arising from the eye movements. Nevertheless, as participants laid still with the head and body fixed, our experiments left as much space as possible for putative fundamental spatiotopic processes to occur (see also^14^). Our results thus rule out space constancy solutions relying on an early encoding of visual space in craniotopic or spatiotopic maps^11,12^.

Moving up the visual hierarchy, responses become more spatially invariant, abstracting away from precise retinal origins. This invariance allows modulation by attention^26,27,49^, multisensory cues, and behavioral demands, but our findings indicate that it does not constitute a shift to spatiotopically organized information. Rather, it is more likely that sources^26,27,49^, such as self-motion^50–52^, pictorial cues^53^, and navigational affordances^54–56^ allow the brain to flexibly direct attention and behaviors to outside-world locations. Interestingly, hippocampus was shown to encode locations in visual space in both humans and rodents^57–59^, indicating that world-centered and sensory-based spatial reference frames can coexist and possibly directly interact at higher levels of processing. Oculomotor behavior is likely central in linking these levels, providing the temporal bridge between sensory input and allocentric spatial maps^60^.

In summary, our study shows that human visual cortex encodes space strictly in retinotopic coordinates, regardless of gaze or attentional focus. By ruling out spatiotopic encoding in visual maps, we shift the focus of space constancy research toward transient motor–sensory interactions and integrative processes across cortical systems. Stable perception emerges not from static world-centered maps in visual cortex, but from the continuous interplay between retinotopic representations, action signals, and higher-level spatial frameworks.

## Methods

### Ethics statement

This experiment was approved by the Ethics Committee of Vrije Universiteit Amsterdam and conducted in accordance with the Declaration of Helsinki. All participants gave written informed consent.

### Participants

Eight students and staff members of the Spinoza Centre for Neuroimaging participated in the experiment (ages 21-40, 1 female, 4 authors). All except two authors were naïve to the purpose of the study and all had normal or corrected-to-normal vision.

### MRI data acquisition

Four T1-weighted (1.0 mm isotropic resolution) and one T2-weighted (1.0 mm isotropic resolution) structural scans were acquired for each participant at The Spinoza Centre for Neuroimaging on a Philips Achieva 3T scanner (Philips Medical Systems, Best, Netherlands) and a 32 channel receive-coil array with a single channel transmit coil. Functional data were collected at the same center on a Philips Achieva 7T scanner (Philips Medical Systems, Best, Netherlands) with a 32 channel receive-coil array with 8-channel transmit coil (Nova Medical, Wilmington, MA). Functional data were collected using a 3D-EPI sequence at a resolution of 1.8 mm isotropic with a 1.3 s volume acquisition time, 44 ms TR, 17 ms TE, consisting of 98 slices of 112 by 112 voxels covering the entire brain. SENSE acceleration was applied in both the Anterior-Posterior (2.61-fold) and Right-Left (3.27-fold) directions. To estimate and correct susceptibility-induced distortions, we acquired an identical EPI image with an opposite phase encoding direction, after each functional scan. Transmit field homogeneity was improved by adjusting the 8-channel transmit output based on a B1+ field population template (i.e., “universal pulse”^61^).

### Stimuli and tasks

The experiment consisted of 3 experimental sessions on different days. Participants started with two fMRI sessions each composed of 10 consecutive runs together with different preparatory and field inhomogeneity mapping scans (about 1h each). The last session was used to obtain multiple structural images (about 45 min in total). Participants were trained on the behavioral task outside the scanner.

Stimuli were presented at a viewing distance of 225 cm, on a 32-inch LCD screen (BOLDscreen, Cambridge Research Systems, Rochester, UK) situated at the end of the bore (17.8 dva horizontally by 10 dva vertically) and viewed through a mirror. The screen had a spatial resolution of 1920 by 1080 pixels and a refresh rate of 120 Hz. Button responses were collected using an MRI compatible button box (Current Designs, Philadelphia, PA, USA). The experimental software controlling the display and the response collection as well as eye tracking was implemented in Matlab (The MathWorks, Natick, MA, USA) using the Psychophysics Toolbox^62,63^.

Participants were instructed to continuously fixate a white bull’s-eye on a black background. The bull’s-eye was composed of a central white dot of 0.04 dva radius surrounded by a white annulus of 0.4 dva radius and a line width of 0.04 dva. They were instructed to attend visual noise contained either within this central bull’s-eye (*attend-fix* condition, 1^st^, 3^rd^, 5^th^, 7^th^ and 9^th^ run) or within a moving bar (*attend-bar* condition, 2^nd^, 4^th^, 6^th^, 8^th^ and 10^th^ run). Each functional session started with 4 runs in which the fixation bull’s-eye was displayed at the screen center together with the moving bar traversing the entire screen (*full-screen* condition, 1^st^, 2^nd^, 3^rd^ and 4^th^ run). In the next 6 runs, the moving bar stimulus was displayed within a 4 dva radius aperture (0.04 dva width cosine edge between the visual noise and the background) surrounding the fixation bull’s-eye. The bull’s-eye was displayed either 4 dva to the left of the screen center (*gaze left* condition, 5^th^ and 6^th^ run), 4 dva to the right of the screen center (*gaze right* condition. 7^th^ and 8^th^ run) or at the screen center (*gaze center* condition, 9^th^ and 10^th^ run). *Full screen* runs lasted 195 seconds (150 TRs). They were composed of 9 interleaved periods with and without the presentation of visual noise. In periods without visual noise, the bull’s-eye was shown alone for 13 seconds (10 TRs). In periods with the visual noise, the bar aperture movement direction was sequentially 180° (left), 270° (down), 0° (right) and 90° (up). The bar aperture could either be horizontally or vertically oriented, in order to be perpendicular to its movement direction. Its center moved in the bar direction on every TR by discrete steps of 0.56 dva, such that 18 vertical and 32 horizontal steps were necessary to traverse the entire screen. *Gaze left, gaze right* and *gaze center* runs lasted 158.6 seconds (122 TRs). They were composed of the same nine interleaved periods with and without the presentation of visual noise. Contrary to the *full screen* runs, the bar aperture movement direction was sequentially 180° (left), 0° (right), 180° (left) and 0° (right) and the bar was always vertically oriented. Its center moved in the bar direction on every TR by 18 discrete steps of 0.22 dva to traverse the circular aperture centered on the bull’s-eye. Visual noise contained within the bar aperture (1 dva width, with 0.04 dva width cosine lateral edges) and the bull’s-eye was composed of distinct randomly generated pink noise (1/f) grayscale textures^64^ (with individual pixel of 0.04 dva width). To probe visual attention, we filtered the orientation contained within the textures to generate clockwise and counterclockwise signal textures. We defined 15 different difficulty levels by either keeping randomly generated noise or by filtering contained orientation by a von Misses distribution with standard deviation values between 10^-1^ (large dispersion) to 10^1.5^ (narrow dispersion) centered around +45° or -45° relative to the vertical axis. Each bar presentations started with 400 ms of streams of unfiltered noise textures presented at 10 Hz in both the bull’s-eye and the bar aperture. This period was followed by 600 ms of clockwise or counterclockwise streams of oriented filtered noise, later followed by 300 ms of unfiltered noise streams. The period with oriented filtered noise streams was highlighted to participants with the bull’s-eye central dot displayed in black. The orientation of the filtered noise was selected randomly and independently every TR for both the bull’s-eye and the bar aperture. Participants reported the orientation of filtered noise streams presented within the bar aperture (*attend-bar* condition) or the bull’s-eye (*attend-fix* condition) with left (-45° or counter-clockwise) or right thumb button presses (45° or clockwise). The difficulty of the task was titrated by a staircase procedure following a 3 down 1 up rule adjusting for the attended stream, the orientation dispersion around ±45°. Staircases started at an intermediate difficulty value (level 10) on every run. Participants had until the end of the TR to press a button if no response was registered the corresponding staircase wasn’t modified.

### Anatomical data preprocessing

T1-weighted (T1w) images were corrected for intensity non-uniformity^65^. The T1w-reference was then skull-stripped. Brain tissue segmentation of cerebrospinal fluid (CSF), white-matter (WM) and gray-matter (GM) was performed on the brain-extracted T1w^66^. A T1w-reference map was next computed after registration of 4 T1w^67^. Brain surfaces were reconstructed using freesurfer^68^ and the brain mask estimated previously was next refined^69^, with the aid of the additional T2w acquisition. Finally, the surface reconstruction was manually edited to correct for pial segmentation errors near the boundary of the occipital lobe and the cerebellum.

### Functional data preprocessing

For each of the 20 functional runs per participant (across all tasks and sessions), the following preprocessing was performed. First, a reference volume and its skull-stripped version were generated using a custom methodology of fMRIPrep^70^. Head-motion parameters with respect to the BOLD reference (transformation matrices, and six corresponding rotation and translation parameters) are estimated before any spatiotemporal filtering^71^. A fieldmap was estimated based on two echo-planar imaging (EPI) references with opposing phase-encoding directions^72^. Based on the estimated susceptibility distortion, a corrected EPI (echo-planar imaging) reference was calculated for a more accurate co-registration with the anatomical reference. The BOLD reference was then co-registered to the T1w FreeSurfer^73^. Co-registration was configured with twelve degrees of freedom to account for distortions remaining in the BOLD reference. The BOLD timeseries were resampled onto their original, native space by applying a single, composite transform to correct for head-motion and susceptibility distortions. Low frequency drift up to 0.01 Hz of the functional time series was removed using cosine drift regressors obtained from fMRIPrep. Functional signal units were next converted to z-score.

### Gaze position monitoring

To ensure participants maintained stable fixation throughout each experimental run, we employed the DeepMREye method to estimate gaze coordinates from eye-region voxels in the raw EPI data^19^ . This analysis validated that all participants, who had received thorough task training, successfully maintained steady fixation across gaze conditions. The analysis procedure involved the fine-tuning of a deep neural network model using a leave-one-participant-out cross-validation approach to determine gaze coordinates based on the expected eye positions for each experimental condition. For each participant, we extracted decoded x and y gaze coordinates over time across the *full screen, gaze center, gaze left*, and *gaze right* runs. To quantify fixation stability, we calculated the Euclidean distance between expected (target fixation location) and decoded gaze positions. Results demonstrated that participants consistently maintained their gaze near the intended locations (*full screen*: 0.05° ± 0.01 —mean ± SEM Euclidian distance across participants—; *gaze center*: 0.04° ± 0.01; *gaze left*: 0.05° ± 0.01; *gaze right*: 0.05° ± 0.01).

### Regions of interest definition and pRF parameters

We first averaged the timeseries of the 8 *full screen* runs, irrespective of whether participants attended to the fixation bull’s-eye or to the bar stimulus. We analyzed this *fiducial* averaged time series using an isotropic Gaussian pRF model^15^. The model included a visual design matrix (converting the visual stimulus in an on-off contrast map for each TR), the position (x, y), the size (standard deviation of the Gaussian), as well a signal amplitude and signal baseline as parameters. The time series were fitted in two parts. The first part was a coarse spatial grid search of 40 linear steps in the 3 spatial model parameters: position (x, y), and size. A linear regression between the predicted and measured time series signal was used to determine the baseline and amplitude parameters. The best-fitting parameters of the first step were next used as the starting point of an optimization phase to produce finely tuned estimates. The visual stimulus was down sampled by a factor 8 for the first of these fitting stages. The grid search position and size parameters were distributed linearly within 20 dva from the center of the screen. In the fine search stage, we used a gradient descent algorithm starting from the obtained grid search parameters, leaving the parameter ranges essentially unconstrained. PRF polar angle maps were derived with the best explained pRF position parameters. The polar angle was drawn using *Pycortex*^74^ on an inflated and flattened cortical visualization of individual participant anatomy. The explained variance of the model set as the coefficient of determination (R^2^) was used to determine color transparency. From this analysis we manually defined for each participant 12 cortical visual regions of interests following pRF angle and eccentricity progression on the cortex as well as anatomical references. These regions include V1, V2 and V3, V3AB which unites V3A and V3B subregions, Lateral Occipital area (LO) which unites LO1 and LO2 subregions, Ventral Occipital (VO) including selective voxels of the ventral path, hMT+ which unites both MT and MST subregions, the inferior and superior part of the Intra Parietal Sulcus areas (iIPS and sIPS) and the inferior, superior and medial parts of the Pre Cenral Sulcus areas (iPCS, sPCS and mPCS). For each ROI, we defined the best-fitting voxels parameters as the 250 highest R^2^ per ROI. We evaluated the effect of the task on the explained variance of the pRFs by taking the task-related time series independently. The selection of 250 voxels was determined through preliminary analyses in which we systematically tested the Bayesian decoder (see below) with varying voxel counts (ranging from 50 to 500 voxels) to identify the optimal number; where decoding performance plateaued while maintaining computational tractability. This analysis revealed that performance gains diminished beyond 250 voxels, making this an efficient choice that balanced statistical power with computational efficiency. To ensure consistency across all analyses, we standardized this voxel selection criterion throughout the study, with the same 250 highest R^2^ voxels used across all conditions and participants within each ROI.

### Out-of-sample predictions

We first evaluated the explained variance of retinotopic or spatiotopic pRF models by applying the parameters obtained in the *full screen* runs to the *gaze* runs time series. To do so, we predicted out-of-set timeseries of the gaze conditions visual designs knowing the pRF parameters of the attention task-specific *full screen* condition to which we either subtracted (*pRFx*_*gaze left*_ = *pRFx*_*full screen*_ *– 4*) or added (*pRFx*_*gaze right*_ = *pRFx*_*full screen*_ *+ 4*) the gaze direction change to the *pRFx* parameter. Signal amplitude (beta) and signal baseline parameters were also adjusted to fit with the new timeseries. Change in explained variance between condition was computed by subtracting the averaged R^2^ observed in the *gaze left* and *gaze right* condition from the R^2^ observed in the *gaze center* condition.

### Fitting predictions

We refine our analysis by determining anew the model parameters in the gaze conditions. To do so, we kept the *pRFy* and *pRFsize* parameters obtained from the *full screen* runs, while fitting anew the *pRFx* parameter as well as amplitude and the baseline. To avoid any bias toward the retinotopic or the spatiotopic model, we started the coarse fitting procedure with *x* coordinate spatial grids centered either on the gaze or on the screen coordinates. The best model (highest R^2^) was next used in the optimization phase to produce finely tuned parameters.

### Bayesian decoding

We developed a novel method to probabilistically decode the position of our moving bar stimulus from the multivariate fMRI data, within a Bayesian framework. The method is based on the standard pRF model fitted on an independent training dataset from the same participant. Provided with a multivariate fMRI timeseries *Y* = *y*_1..*n*,1..*t*_ with *n* voxels and *t* time points, the method yields a posterior distribution *p*(*x*_1..*t*_, *h*_1..*t*_| *Y*) over possible stimulus positions *x* and heights *h* for every frame in the time series where a stimulus was presented. The method allows to probe what bar position is represented on the level of a given ROI, on a TR-to-TR basis.

By jointly sampling across all time points of the data concurrently, we took into account the hemodynamic delay and hemodynamic temporal smoothing in the estimation procedure. This is made computationally feasible by a state-of-the-art computational graph library developed for deep neural network training (Tensorflow), massively parallel graphics processing units for computation (GPUs), as well as Hamiltonian MCMC method^75^, which exploits the exact gradient functions that the “*autodiff*” functionality that Tensorflow provide.

To obtain a posterior distribution over bar positions, given fMRI data, we first needed a likelihood function that describes the probability of the data given a stimulus time series *S*. Let’s assume *S* a time series of 2D stimulus images. The likelihood function is the probability density function of a multivariate *t* distribution over the residuals of the pRF model with a covariance matrix *Σ* of a multivariate t-distribution with *df* degrees-of-freedom. Hence, *Y* = *pRF*(*S*; *θ*_1..*n*_) + *ϵ* where *ϵ* ∼ *t*(0, *Σ, df*) and thus *p*(*Y*|*x*_1..*t*_, *h*_1..*t*_; *θ*_1.. *n*_, *Σ, df*) = *t*(*Y* − *prf*(*S*; *θ*_1..*n*_); *Σ, df*) where *pRF* is the output of a standard pRF model for the image time series *S*, as described in the previous section, with voxelwise position, dispersion, amplitude, and baseline parameters *θ*_1..*n*_ estimated using standard pRF fitting methods on an independent training dataset.

The pRF function includes a convolution step where “neural” time series are convolved with a canonical hemodynamic response function. This means that the pixel intensities on timepoint *t* influence the residuals and thereby likelihoods of multiple frames of neural activity data (roughly between timepoint *t* and 20 seconds later). This also means that different dimensions of the likelihood function that are close together in time are correlated, necessitating more advanced samplers than traditional Gibbs sampling^76,77^.

To estimate the noise covariance matrix Σ, we couldn’t use the sample covariance of the residuals of the training set, because these estimates would be highly unstable in our data regime of sparse data (a few hundred time points) and a large number of dimensions^78^. Therefore, we estimated a regularized covariance matrix 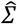, using a method developed for decoding Gabor orientations from fMRI signals in early visual areas^16^. Specifically, the covariance matrix estimate 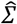 is a weighted sum of a diagonal covariance matrix *I* ∘ *ττ*^*T*^, a perfectly correlated covariance matrix *ττ*^*T*^, and a matrix *WW*^*T*^, where *W* is a *n* × *m* matrix with *n* the number of voxels and *m* the total number of pixels in a single stimulus image. Each element of the matrix *W* describes the linear weight of its respective pixel to the activation of its respective voxel. More specifically, this value is the probability density of the *x* and *y* coordinates for the multivariate Gaussian described by the pRF parameters *θ* for that voxel, multiplied by its estimated amplitude.

Thus, we assume that the covariance of the residuals of two voxels that have particularly similar pRF should generally be higher than that of those voxels of which the pRF do not overlap^16–18^. The complete formula for the estimated residual covariance matrix 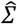 is: 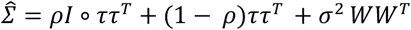 where the vector *τ* > 0, scalar 0 ≤ *ρ* < 1, and scalar σ^2^ ≥ 0 are estimated using maximum likelihood estimation on the residuals *Y*^*training*^ − *pRF*(*S*^*training*^; *θ*_1..*n*_) of the training data, using gradient descent optimization. However, the dimensionality of the stimulus time series *S* was so high (above 100,000) that it was infeasible to sample its posterior distribution. Moreover, its dimensions are not semantically meaningful. Therefore, we opted to drastically reduce the dimensionality of *S* using a two-parameter stimulus model *s*(*x, h*) that returns a two-dimensional pixel image with a vertically centered bar of a fixed width, centered on the *x* coordinate and a height of *h*. We chose this stimulus model because both the x position and height of the bar varied over time in the gaze conditions. Note that, in the stimulus function, the pixel intensity rapidly but smoothly falls off as a sigmoidal function of distance to the border of the bar stimulus. Such a smooth functional form was necessary to keep the likelihood function differentiable and thereby Hammoltonian MCMC feasible. This way, the number of parameters of the likelihood function was heavily reduced to 2 times the number of timepoints: *p*(*Y*| *x*_1..*t*_, *h*_1..*t*_; *θ*) = *t*(*Y* − *pRF*(*s*(*x*_1..*t*_, *h*_1..*t*_); *θ*_1..*n*_); Σ, *df*). We reduced this number a bit further by not estimating stimulus properties for frames where there was no stimulus bar on the screen such that the model assumed these frames to be an empty screen with a fixation bull’s-eye.

We used uniform priors on the estimated *x* coordinates, restricted between the leftmost and rightmost coordinates on the screen as well as uniform priors on the height of the stimulus bar between 0 and the height of the stimulus screen. This was achieved using bijective sigmoidal transformation functions. The entire procedure was implemented in TensorFlow 2 and packaged as a freely available Python package called Braincoder (https://github.com/Gilles86/braincoder).

As a first step, a 2D grid search (position by height) was performed for every time point separately, optimizing the likelihood function with respect to single frame *x* and *h* parameters, neglecting the covariance structure of the likelihood function. After this grid search, a maximum a posteriori (MAP) estimate for the entire time series *argmax p*(*x*_1..*t*_, *h*_1..*t*_|*Y*) was estimated using gradient descent, starting from the values found in the grid search. Finally, this MAP estimate was used as a starting point for the NUTS MCMC sampler, a self-tuning Hamiltonian MCMC sampler^77^, to obtain samples from the posterior *p*(*x*_1..*t*_, *h*_1..*t*_|*Y*). We used 4 independent chains, with 250 steps and 500 samples per chain. After some experimenting, we settled on an acceptance probability of 0.3 to more effectively sample our highly complex 144-dimensional posterior distribution.

### Reference frame index

We devised a reference frame index (RFI), based on earlier work on the topic^11,13^. This index contrasts the ability of a spatiotopic versus a retinotopic model to explain the observed data. Specifically, it expresses the difference in explained variance between the retinotopic and the spatiotopic model as a fraction of the sum of both explained variances. Hence, the index ranges between -1 (completely and exclusively consistent with spatiotiotopic reference frame) and 1 (completely and exclusively consistent with the retinotopic reference frame):

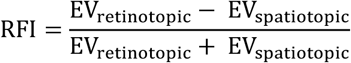

Where EV is the reduction in variance from the raw data versus the residuals of the model fit EV = Var(Y − Ŷ) − Var(Y). Note that for the out-of-sample predictions (Fig. 3) of individual PRF time series, the variance is defined on real and predicted BOLD timeseries of individual voxels. For the decoding analysis (Fig. 4) the variance is defined on the actual and the decoded spatial positions of the bar stimulus.

### Statistics

For statistical comparisons we used SciPy^79^ paired permutation test with 10,000 permutation bootstrapped iterations used to approximate the null distribution^80^. This procedure allows us to resample our data to create a distribution with randomly rearranged labels of the conditions to compare for each participant. We determined statistical significance by deriving two-tailed (see Results) *p* values for the comparison of the distributions of the compared conditions.

### Voxel reliability and spatial relevance analysis

To investigate the relationship between reference frame indices and voxel characteristics that affect measurement reliability, we conducted a comprehensive analysis examining all voxels within each ROI. This analysis was designed to address two key factors that influence the informativeness of individual voxels for distinguishing between retinotopic and spatiotopic hypotheses. First, we quantified voxel reliability using the explained variance (R^2^) obtained from the pRF model fit in the *full screen* runs. Voxels with higher explained variance represent more reliable measurements that are less influenced by estimation noise and overfitting, and therefore provide more robust estimates for subsequent model comparisons. Second, we calculated the spatial relevance of each voxel by determining the percentage of overlap between the voxel’s pRF and the central 4 dva radius aperture used in the *gaze* conditions. This overlap percentage was computed by calculating the intersection area between the voxel’s fitted Gaussian pRF (defined by its position and size parameters from the *full screen* runs analysis) and the circular aperture centered on the fixation bull’s-eye. Voxels with higher overlap percentages are more informative for distinguishing between reference frames, because both retinotopic and spatiotopic models make clear and distinguishable predictions for these voxels across the different gaze conditions. Conversely, voxels with minimal overlap (particularly those at extreme eccentricities or outside the aperture) would show unreliable responses under the retinotopic hypothesis and reduced reliability in most conditions under the spatiotopic hypothesis.

For this analysis, we computed reference frame indices for all voxels within each ROI using the out-of-sample prediction approach described previously. The RFI was calculated by comparing the explained variance of strictly retinotopic versus strictly spatiotopic pRF models when predicting timeseries from the *gaze left* and *gaze right* runs using parameters derived from the *full screen* runs. We then created two-dimensional plots for each ROI, with explained variance on one axis and percentage pRF-aperture overlap on the other axis, with each voxel color-coded according to its reference frame index. This visualization approach allowed us to examine how reference frame preferences relate to both the reliability (explained variance) and spatial relevance (aperture overlap) of individual voxels, providing insight into whether any spatiotopic tendencies might be driven by less reliable or spatially irrelevant measurements.

## Data and code availability

We make available online our imaging dataset, including all derivatives and individual participant and group figures (openneuro.org/datasets/ds004091). Brain models with data analysis visualization are made available online (invibe.nohost.me/gazeprf) together with the experimental and data analyses codes (github.com/mszinte/gaze_prf).

## Acknowledgments

We are grateful to the members of the Knapen and Dumoulin laboratories in Amsterdam for helpful comments and discussions and to Alice and Clémence Szinte and Candice Duval for their support. Centre de Calcul Intensif d’Aix-Marseille is acknowledged for granting access to its high-performance computing resources. This research was supported by a Marie Skłodowska-Curie Action Individual Fellowship (704537) and a Agence Nationale de la Recherche JCJC (RetinoMaps) grant to M.S. as well as an NWO-CAS (012.200.012) and ABMP (2015-7) grants to T.K.

## Author contributions

Conceptualization, M.S., and T.K.; Methodology, M.S., G.H., M.A., and T.K.; Software, M.S., G.H., M.A., and T.K.;Validation, M.S., G.H., M.A., and T.K.; Formal Analysis, M.S., G.H., M.A., and T.K.; Investigation, M.S., I.V., and T.K.; Resources, T.K.; Data Curation, M.S.; Writing – Original Draft, M.S., G.H., and T.K.; Writing – Review & Editing, M.S., G.H., M.A., I.V., S.D., and T.K.; Visualization, M.S., G.H. and T.K.; Supervision, S.D., T.K.; Project Administration, T.K.; Funding Acquisition, M.S. and T.K.

---

**Supp. Figure 1.**
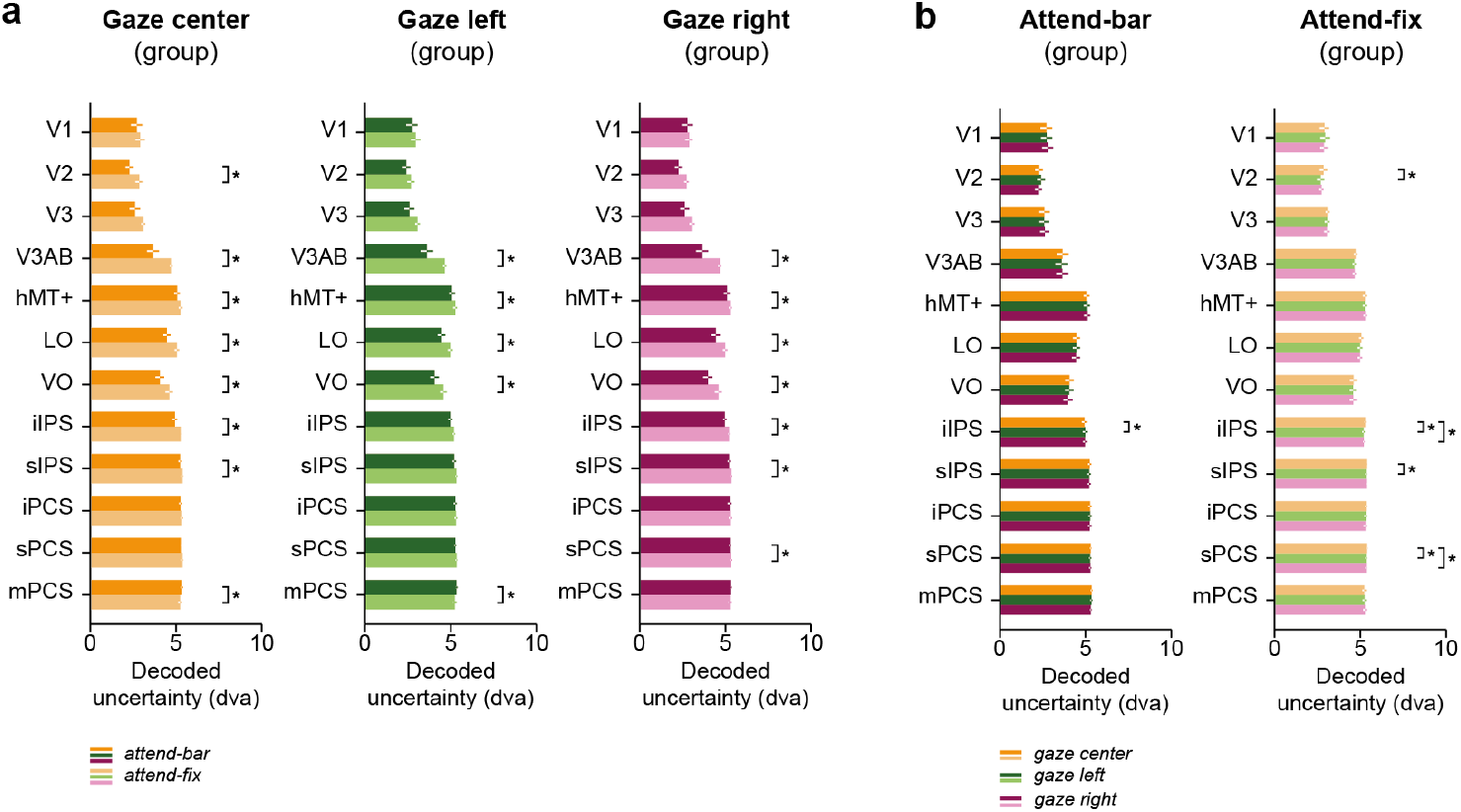
Decoding uncertainty. **a**. Average decoded uncertainty across ROIs and participants for the gaze center (left column), gaze left (middle column) and gaze right (right column) conditions. Black asterisks indicate a significant difference between the *attend-bar* and *attend-fix* conditions (two-sided *p* < 0.05). **b**. Same results for the attend-bar (left column) and attend-fix (right column) conditions. Black asterisks indicate a significant difference between the *gaze center, gaze left* and *gaze right* conditions (two-sided *p* < 0.05).

